# Migratory patterns of two major influenza virus host species on tropical islands

**DOI:** 10.1101/2023.01.18.524666

**Authors:** Camille Lebarbenchon, Solenn Boucher, Chris Feare, Muriel Dietrich, Christine Larose, Laurence Humeau, Matthieu Le Corre, Audrey Jaeger

**Affiliations:** Université de La Réunion, UMR Processus infectieux en milieu insulaire tropical (PIMIT), INSERM 1187, CNRS 9192, IRD 249, 2 rue Maxime Rivière, Sainte-Clotilde, La Réunion, France; Université de la Réunion, UMR Ecologie marine tropicale des océans Pacifique et Indien (ENTROPIE), CNRS IRD, IFREMER, Université de Nouvelle-Calédonie, 15 Avenue René Cassin, Saint Denis, La Réunion, France; WildWings Bird Management, Haslemere, Surrey, United Kingdom; Université de La Réunion, UMR Peuplements végétaux et bioagresseurs en milieu tropical (PVBMT), CIRAD, 15 Avenue René Cassin, Saint Denis, La Réunion, France

**Keywords:** Brown noddy, Lesser Noddy, Avian influenza, Tracking, Serology, Seychelles

## Abstract

Animal migration is a major driver of infectious agent dispersal. Duck and seabirds migrations, for instance, play a key role in the spatial transmission dynamics and gene flows of avian influenza viruses (AIV), worldwide. On tropical islands, brown and lesser noddies (*Anous stolidus* and *A. tenuirostris*) may be important AIV hosts, but the lack of knowledge on their migratory behaviour limits our understanding of virus circulation in island networks. Here we show that high connectivity between islands generated by post-breeding behaviours could be a major driver in the spread and the maintenance of AIV among tropical islands. Tracking data highlight two types of behaviours during the non-breeding season: birds either staying in the vicinity of their breeding ground (on Bird Island, Seychelles), or moving to and roosting on other islands in the Indian Ocean. Differences in roosting places are detected between migrant birds, ranging from the Tanzanian coast to the Maldives archipelago and Tromelin Island. Epidemiological data further support that brown and lesser noddies are major hosts for AIV, although significant variation of seroprevalence between species suggest that other drivers are involved in virus infection and transmission dynamics.

## 1. Introduction

Animal migration is a key mechanism for the dispersal of infectious agents over long distances [1] and could therefore play a significant role in the spread of viruses to and among island ecosystems. Tropical oceanic islands might for instance be connected to the global avian influenza virus (AIV; *Alphainfluenzavirus*) epidemiology [2], but the frequency of virus exchanges between islands and from/to neighbouring continents remains to be precisely assessed. Wild ducks and seabirds are natural hosts for avian influenza virus [3,4]; in particular, seabirds are reservoir for the H13 and H16 virus subtypes [5–7]. In seabirds, AIV have been primarily isolated from Laridae (*e*.*g*. gulls, terns) and Scolopacidae (*e*.*g*. waders). Low prevalences of infected birds are usually reported [8–11], although high temporal variation is likely to characterize AIV transmission dynamics in these hosts, in particular during the breeding season [12]. However, most investigations have focused on AIV transmission on continental habitats [13,14], and virus transmission dynamics and diversity in seabird populations on tropical oceanic islands is yet to be characterised.

The absence of wild ducks on most tropical islands could lead to major differences in AIV ecology and epidemiology, as compared to continental habitats. High seabird breeding-site fidelity [15] could restrict virus dispersal between populations breeding on different islands. Because of the discrete geographical nature of oceanic islands, host population structure could also have major effects on virus transmission between islands as well as on the diversity of viruses circulating on each island. Spatial isolation may create opportunity for the maintenance of viruses in wild bird communities inhabiting these islands, leading to the endemic circulation of certain AIV subtypes and genotypes (*e*.*g*. H15; [16]). Extensive seabird post-breeding migrations may, however, counterbalance this effect by increasing virus dispersal between islands and homogenizing virus diversity between islands.

With an estimated breeding population size of 19 million individuals [17], seabirds are the most abundant and widespread avifauna in the Western Indian Ocean [18]. Small oceanic islands in this region are major breeding sites for terns, with several species aggregating at very high densities in breeding colonies, involving hundreds of thousands, occasionally millions, of birds [19]. In a previous study, we identified the host range of AIV in seabirds in the islands of the Western Indian Ocean, and further assessed the virus subtype diversity based on serological assays [2]. These findings suggested that terns may represent a major and neglected host on tropical oceanic islands. For instance, high prevalence of infection was estimated in Lesser noddy (*Anous tenuirostris*) on Re-union Island (up to 28% of birds shedding virus and 79% of seropositive birds). Such high prevalences contrast with the low level of AIV detection usually reported for terns [13], and are comparable to prevalences usually reported in ducks and gulls [3,20]. Virus gene flow between Eurasia and the Western Indian Ocean was also demonstrated [2], highlighting that in spite of their spatial isolation, tropical oceanic islands are connected to the global AIV epidemiology, and that tern migrations and behaviour may create opportunities for the maintenance of viruses in wild bird communities inhabiting these islands.

In this study, we investigated the AIV dispersive behaviours of two seabirds species (Lesser noddy and Brown noddy, *Anous stolidus*). Based on tracking data, we characterized their post-breeding migration and activity in the Western Indian Ocean. Serology and molecular detection were carried out to assess the prevalence of birds with AIV antibodies and shedding viruses, respectively. Our results highlight the high connectivity between islands, generated by post-breeding migrations, and the key role of Lesser and Brown noddies in the spread of AIV among tropical islands.

## 2. Materials and Methods

### (a) Study site

The study was conducted on Bird Island, the northernmost island of the Seychelles archipelago (3°43’ S, 55°12’ E). Bird Island is a 90 ha low-lying coral sand cay and a major breeding site for terns. Approximately 400 000 Sooty terns (ST; *Onychoprion fuscatus*) pairs nest annually on the ground in the northern part of the island [21]. The colony is a tourist attraction for visitors to the small hotel located in the southern part of the island.

Woodland areas located in the centre of the island provides habitat for three tree-nesting seabird species: the Brown noddy (BN), the Lesser noddy (LN), and the White tern (*Gygis alba*). During the breeding season (April to September), approximately 10 000 BN and 300 LN breeding pairs inhabit the island [21,22]. BN are mainly located in coconut trees and *Casuarina equisetifolia*, with some in decorative *Cordia subcordata* around the hotel and also in low *Scaevola taccada* around the coast and fringing the airstrip. BN also nest on the ground around the hotel. LN nest in the woodland on the western side of the island, mainly in *Pisonia grandis*, but also in *Casuarina equisetifolia* and, around the hotel, in *Cordia subcordata*. White Terns mainly in Casuarinas, but some in smaller trees and bushes.

### (b) Research permits and ethic statements

Field work and collection of biological material in the Seychelles were approved by the Seychelles Bureau of Standards and the Seychelles Ministry of Agriculture, Climate Change and Environment. Birds capture, handling, and marking was also approved by the Center for Research on Bird Population Biology (National Museum of Natural History, Paris; Program 616) and the British Trust for Ornithology. All procedures were evaluated by an ethic committee (Comité d’ éthique de La Réunion; agreement number A974001) and authorized by the French Ministry of Education and Research (APAFIS#3719-2016012110233597v2).

### (c) Seabird tracking

BN and LN at-sea distribution and activity were investigated with Global Location Sensors (GLS). GLS were attached to stainless steel leg rings with cable ties, on the tarsus of incubating adults. The combined mass of logger, leg band, and cable ties represented less than 3% of the body mass and thus was within acceptable mass limits for devices attached to seabirds [23]. In 2012, 25 GLS (MK18 model, British Antarctic Survey, United Kingdom) were deployed on BN as a preliminary study. Fifteen (60%) were recovered the following year (data were successfully downloaded from all the 15 GLS). Given the recovery rate, additional GLS were then deployed in 2014, on BN (17 MK3006, Biotrack Ltd., United Kingdom), but also on LN (17 MK5093, Biotrack Ltd., United Kingdom). Fifteen GLS were recovered from this second set of equipped birds, and data were successfully downloaded from 12 GLS.

GLS devices record elapsed time and light level, allowing estimates of geographic position twice per day with an average spatial accuracy of 186 km for birds in flight [24]. Bird locations were estimated using the threshold-method with the geolight package [25]. We then removed unrealistic positions yielding unrealistic flight speed [26]. Fifty percent kernel density distributions (core areas) were calculated to examine post-breeding noddy at-sea distribution with the adehabitat package [27]. The date and length of migrations and long foraging trips, were determined by identifying rapid shifts in distance from the colony for each bird. We identified roosting places by identifying the closest island to the centroid of each post-breeding individual core area.

GLS also test for saltwater immersion every 3 seconds and record number of positive tests from 0 (continuously dry) to 200 (continuously wet) each 10 minutes. We thus also estimated the percent of time spent in contact with seawater [28], during day and night (categorized with light data), and for both the breeding and post-breeding periods. The fixed effects of sex, species, photoperiod (day and night data) and season (breeding and post-breeding data) on the percentage of time spent on water were investigated using a GLMM with individuals as random effect. Analyses were performed with R 3.4.2 [29].

### (d) Bird sampling

Samples were collected from BN, LN, and also ST (details available in supplementary table 1), in order to investigate species-related and temporal variation in AIV shedding and antibodies between the three most abundant tern species on Bird Island. For each bird, faeces (cloacal swab) and saliva (oropharyngeal swab) were collected with sterile rayon-tipped applicators (Puritan, Guilford, ME, USA). Both swabs were placed in a single tube, containing 1.5 ml of Brain Heart Infusion (BHI) media (Conda, Madrid, Spain) supplemented with penicillin G (1000 units/ml), streptomycin (1 mg/ml), kanamycin (0.5 mg/ml), gentamicin (0.25 mg/ml), and amphotericin B (0.025 mg/ml). Swabs were maintained at 4°C in the field, shipped to the laboratory within 48 hours, and held at −80°C until tested. A small sample of blood (up to 1.0% of body weight) was collected from the medial metatarsal vein and centrifuged within 4 hours after collection. Sera were transferred in cryotubes and stored at −20°C. Samples were shipped to the laboratory within 48 hours and held at −20°C until tested.

### (e) Serology

Sera were tested with the IDvet ID Screen Influenza A Antibody Competition (IDvet, Montpellier, France) enzyme-linked immunosorbent assay (ELISA), following an optimized protocol for the detection of antibodies specific to the AIV nucleoprotein (NP) in wild birds [30]. This protocol was also used for the detection of seropositive seabirds on Bird Island, in a previous study [2]. Sample absorbance was measured at 450 nm with a Sunrise microplate reader (TECAN, Grödig, Austria). Samples with a sample-to-negative control ratio (S/N) below 0.4 were considered positive for the presence of AIV NP antibodies; samples with S/N greater than or equal to 0.55 were considered negative. Samples that yielded S/N between 0.4 and 0.55 were re-tested and, following the S/N obtained in the second test, were considered either negative (S/N > 0.4) or positive (S/N < 0.4).

Previously published data were included in the statistical analysis (N = 568 sera collected in 2012 and 2013; supplementary table 1). Chi square tests (χ^2^) were performed to test the effect of the bird species (BN, LN, ST) and the breeding season (2013, 2013, 2014, 2015) on the probability of successful detection AIV NP antibodies in bird serum, with R 3.4.2 [29]. Given the low number of sampled chicks (N = 33) and juveniles (N = 4), only adult birds were included in the statistical analysis (N = 1246; supplementary table 2)

### (f) Molecular detection

Samples (swabs) were thawed overnight at 4°C, briefly vortexed and centrifuged at 1 500 *g* for 15 min. RNA extraction was performed with the QIAamp Viral RNA Mini Kit (QIAGEN, Valencia, CA, USA). Reverse-transcription was performed on 10 µL of RNA, with the ProtoScript II Reverse Transcriptase and Random Primer 6 (New England BioLabs, Ipswich, Massachusetts, USA), under the following thermal conditions: 70°C for 5 min, 25°C for 10 min, 42°C for 50 min and 65°C for 20 min. cDNA were tested for the presence of the AIV Matrix (M) gene by real-time polymerase chain reaction (rt-PCR) [31] with a CFX96 Touch Real-Time PCR Detection System (Bio-Rad, Hercules, CA, USA).

### (g) Molecular sexing

Molecular sexing was performed for birds equipped with GLS [32]. DNA was extracted from blood samples with the QIAmp Blood & Tissue kit (QIAGEN, Valencia, CA, USA). PCR reactions were performed with 7.5 μL of GoTaq G2 Hot Start Green Master Mix (Promega, Madison, WI, USA), 0.6 μL of each primer (10 μM) and 50 ng of DNA template (4 μL), in a final volume of 15 µL. Amplifications were performed using a GeneAmp PCR System 9700 (Thermo Fisher Scientific, Waltham, MA, USA) under the following thermal conditions: 94°C for 2 min, followed by 40 cycles at 94°C for 30 s, 50°C for 30 s, and 72°C for 45 s, and by a final elongation at 72°C for 4 min. PCR products were size-fractioned in a 1.5% agarose gel stained with GelRed nucleic acid gel stain (FluoProbes).

## Results

### (a) Post-breeding at-sea distribution

Two behaviours were highlighted in BN and LN during the post-breeding period: birds remaining in the vicinity of their breeding ground (residents: BN = 8, LN = 2) and birds moving to and roosting on another island (migrants: BN = 12, LN = 5) (figures 1 and 2). Proportions of residents and migrants were not statistically different between species (Fisher’ s exact test, p = 0.678), and between females and males. Indeed, for BN, 80% of females and 40% of males were migrants (Fisher’ s exact test, p = 0.17), and for LN, 75% of females and 67% of males were migrants (Fisher’ s exact test, p = 1).

**Figure 1.**
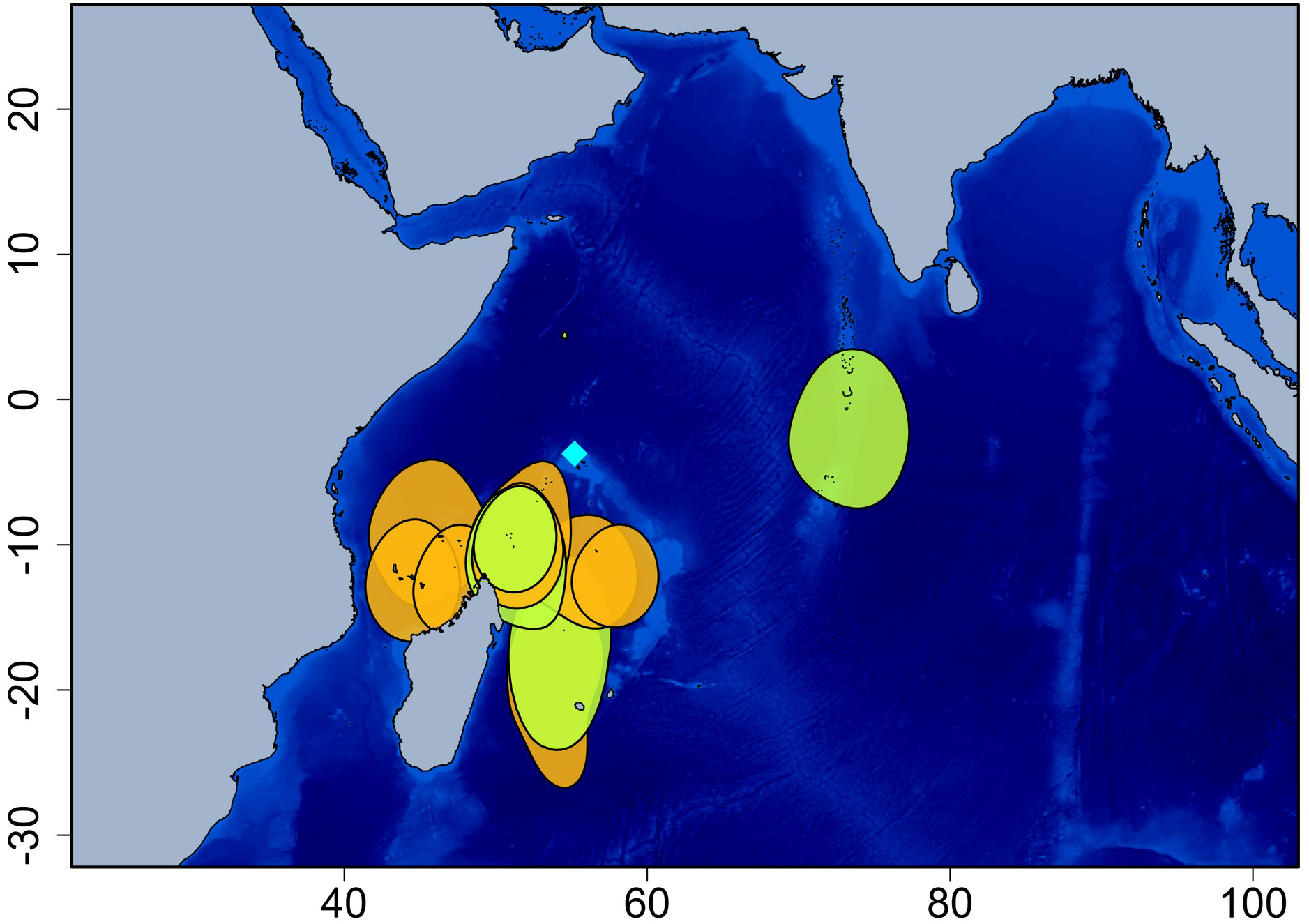
Brown noddy (*Anous stolidus*) 50% kernel density distribution (migrant birds). The blue diamond indicates Bird Island (breeding colony) and black dots indicate roosting islands (i.e. closest island to the centroid of each post-breeding individual core area). Orange: females; Green: males.

**Figure 2.**
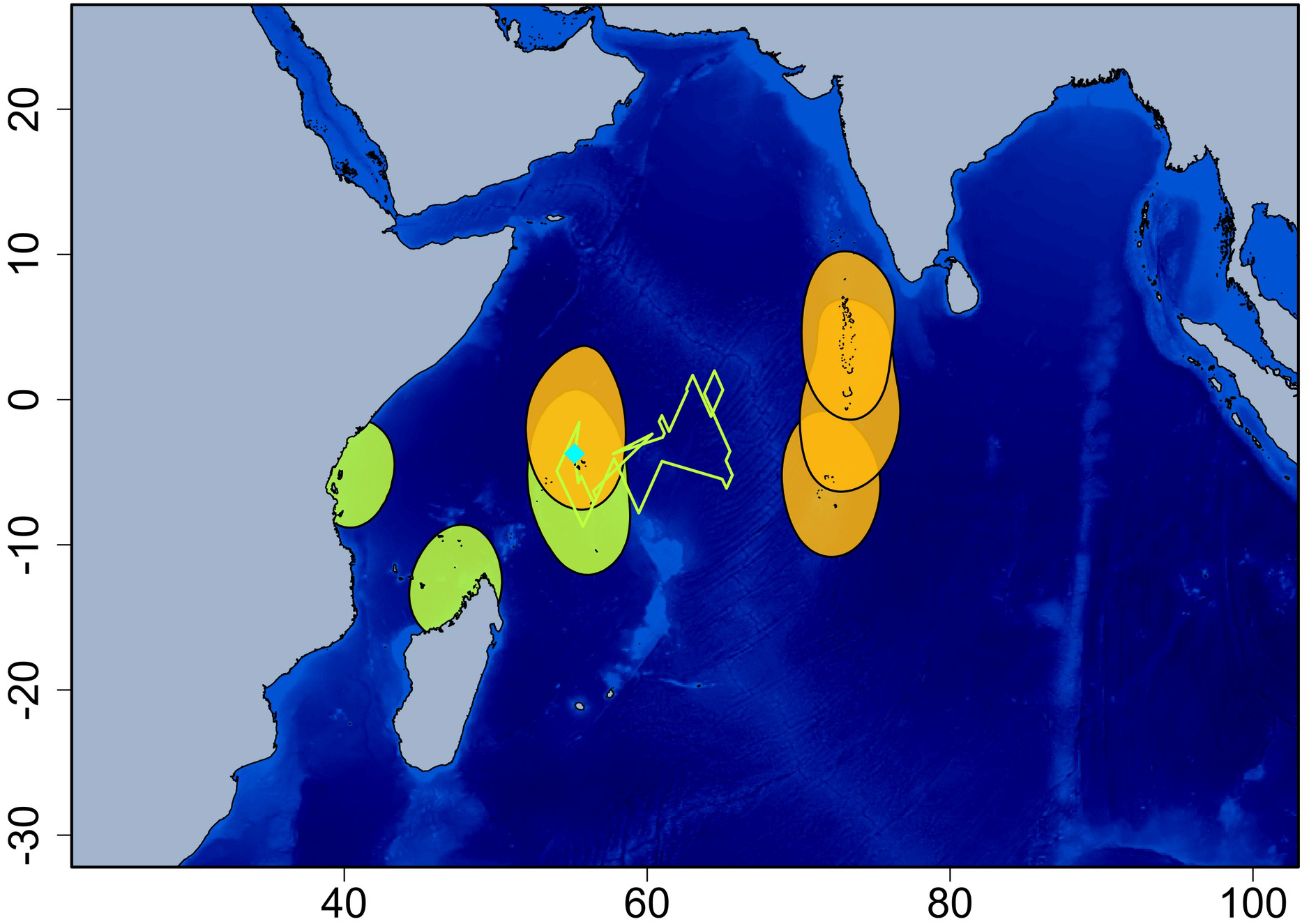
Lesser noddy (*Anous tenuirostris*) 50% kernel density distribution (resident and migrant birds). The blue diamond indicates Bird Island (breeding colony) and black dots indicate roosting islands (i.e. closest island to the centroid of each post-breeding individual core area). The green track indicates a long foraging trip conducted by one resident bird. Orange: females; Green: males.

Migrant birds left the colony for 159.1 ± 35.3 and 174.5 ± 52.6 days for BN and LN, respectively (Mann-Whitney test, p = 0.489). All migrants used one specific roosting place except one BN individual that visited two different places during the post-breeding period. Migrant BN roosted on seven different places, mostly to the south of the Seychelles, on Tromelin Island (N = 2), Farquhar Group (N = 5), Agalega Island (N = 2), Aldabra Islands (N = 1), Comoro Islands (N = 1), and in the north-west coast of Madagascar (N = 1); except one bird roosting to the East on the Maldives archipelago (figure 1). For LN, migration direction differed between males and females: all three migrant females roosted East of the Seychelles, on the Maldives (N = 2) and Chagos (N = 1) archipelagos, while both migrant males roosted to the West of the Seychelles on the Tanzanian coast (N = 1) and the north-west coast of Madagascar (N = 1) (figure 2).

Migrant BN, but not LN, performed long foraging trips from their roosting places (number of foraging trips per individual: 0.8 ± 1.0; average distance: 1 820.4 ± 1 075.3 km; average duration: 33.6 ± 23.2 days). Resident BN also performed long foraging trips (2.5 ± 0.8 per individuals; 2 244.2 ± 977.8 km; 64.4 ± 51.1 days). For LN, one resident bird performed a long foraging trip during 17 days and 1 206 km distant from the colony (figure 2, green track).

### (b) At-sea activity

The proportion of time spent on water varied daily and seasonally (GLMM, F = 287.5 and 189.1, respectively, both p < 0.0001), but no variation was observed between females and males (GLMM, F = 0.6, p = 0.433) nor between species (GLMM, F = 3.6, p = 0.072) (figure 3). Indeed, birds spent more time on water during the day (20.5 ± 10.6 % for BN and 19.7 ± 11.8 % for LN) than during the night (4.7 ± 5.6 % for BN and 1.3 ± 2.0 % for LN), and during the post-breeding period (19.5 ± 12.0 % for BN and 16.5 ± 14.9 % for LN) than during the breeding period (5.6 ± 5.4 % for BN and 4.5 ± 4.9 % for LN).

**Figure 3.**
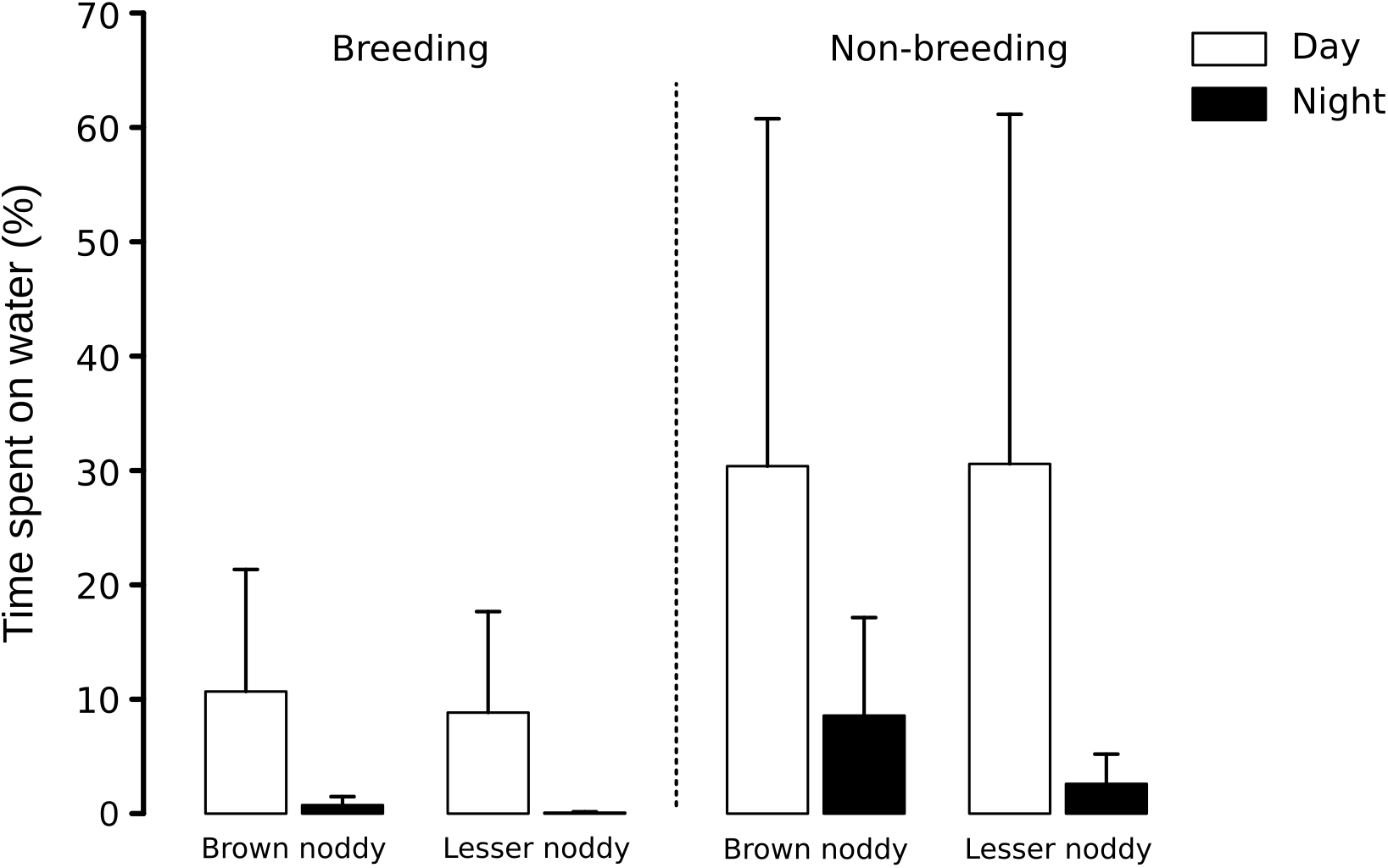
Proportion of time spent in contact with seawater for Brown noddy (*Anous stolidus*) and Lesser noddy (*Anous tenuirostris*).

### (c) Molecular detection and seroprevalence of avian influenza virus

In total, cloacal and oropharyngeal swabs were obtained from 720 birds: 284 BN (N = 151 in 2014 and N = 133 in 2015), 274 LN (N = 173 in 2014 and N = 101 in 2015), and 162 ST (N = 109 in 2014 and N = 53 in 2015). None of the samples tested positive for the presence of the AIV M gene by rt-PCR.

Overall, significant differences in the prevalence of seropositive birds were detected between species (χ^2^ = 241; df = 2; p < 0.001) and between years (χ^2^ = 32; df = 3; p < 0.001) (figure 4). Seroprevalence was higher for LN (58 ± 4.9 %) than for BN (27 ± 4.3 %) and for ST (7.9 ± 2.4 %); it was also higher in 2014 (36 ± 4.7 %) than in 2013 (32 ± 5.5 %), 2015 (31 ± 5.3 %), and 2012 (17 ± 4.3 %). When each species was considered independently, however, inter-annual variations were no longer significant (BN: χ^2^ = 5.3; df = 3; p = 0.15; LN: χ^2^ = 2.1; df = 3; p = 0.55; ST: χ^2^ = 1.1; df = 3; p = 0.77).

**Figure 4.**
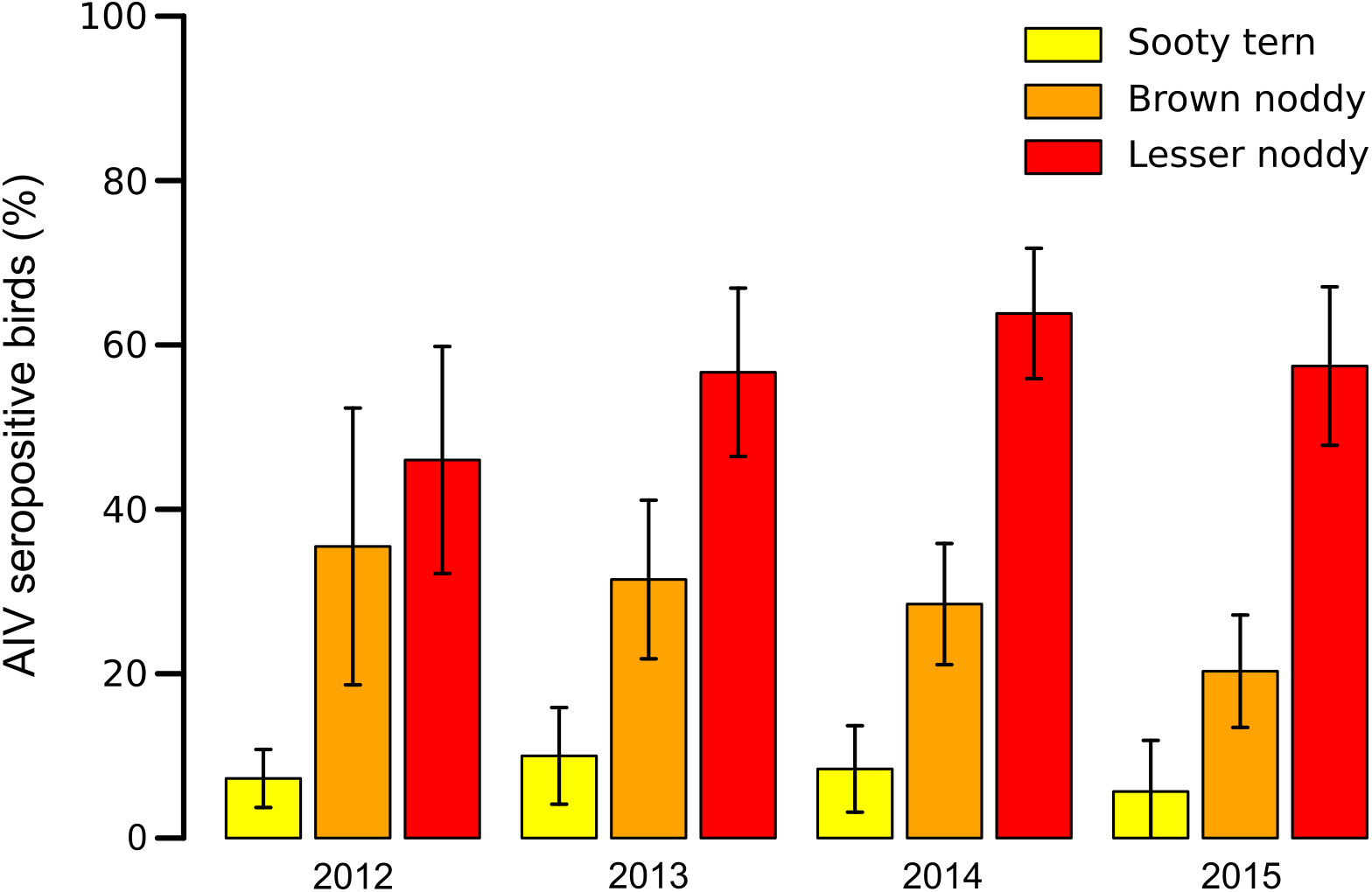
Prevalence of Avian Influenza Virus (AIV) seropositive adult birds (± 95% confidence interval).

Individual variation of AIV NP specific antibodies among years was investigated for recaptured birds. For BN, 63 birds were tested at least twice (for two years), between 2012 and 2015. For LN, only 10 birds sampled in 2014 were recaptured and tested in 2015. Among those birds, 27% and 70% tested positive at least one year, for BN and LN, respectively. For BN, but not for LN, different seroconversion patterns were found (figure 5 and supplementary table 3). Although most seropositive birds remained positive for consecutive years, four birds acquired NP antibodies between the first and the second year (GE54841, GE54824, DE92952 and DE92989), and three birds tested negative after being previously positive (DE92967, DE95924, GE41756). One bird tested negative the first year, positive the following one, and turned seronegative again the third year (GE54837; figure 5).

**Figure 5.**
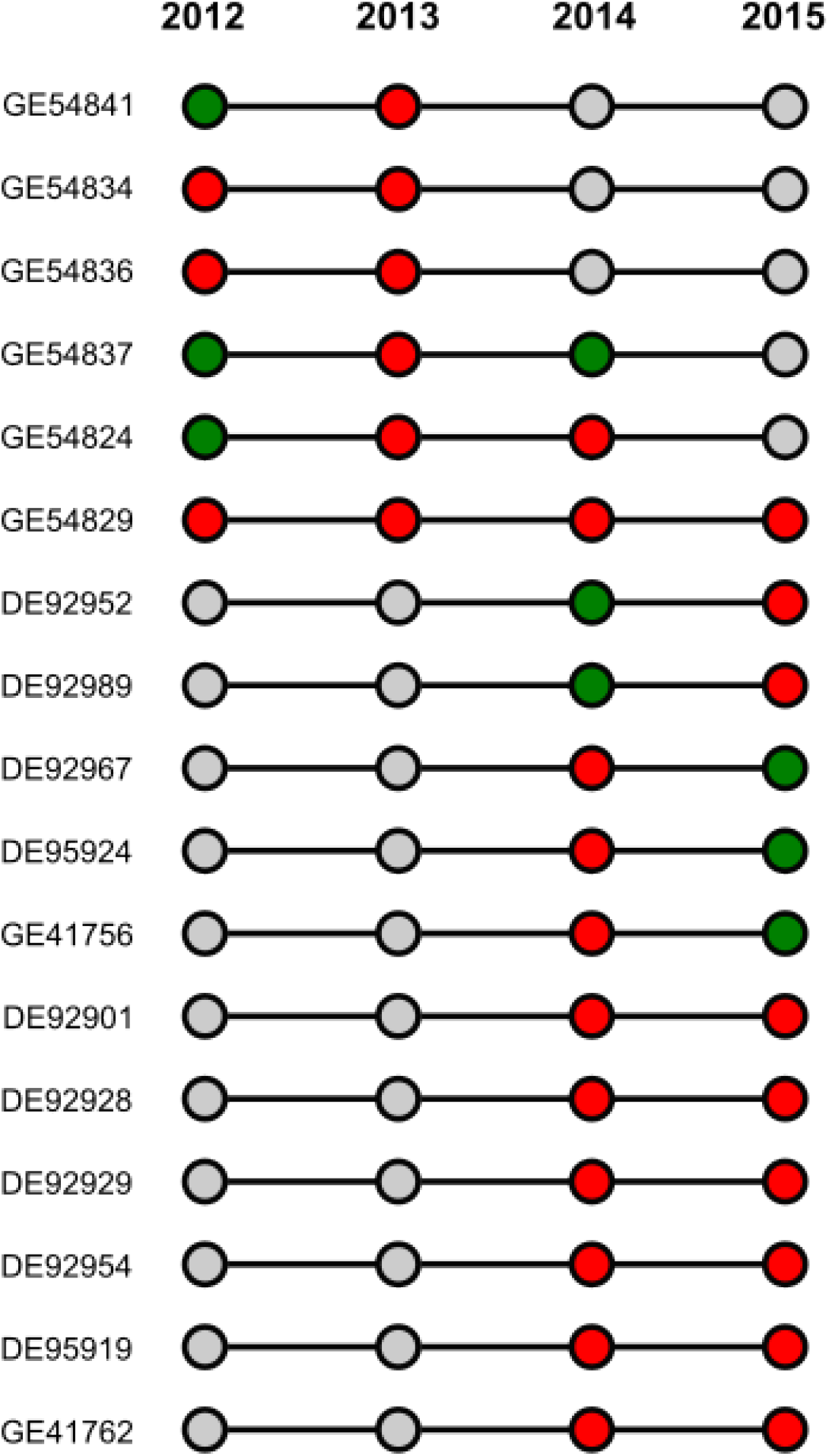
Individual patterns of Influenza A virus nucleoprotein specific antibodies in Brown noddies (*Anous stolidus*). Only birds that tested positive at least one year are included. Green circle: seronegative bird; red circle: seropositive bird; grey circle: not sampled.

## Discussion

Host migrations favour the mixing of AIV between flyways [33] and intercontinental gene flows [34], in particular for seabirds [4,7]. Incorporating epidemiological data and animal movement analyses can provide critical insight for the understanding of virus transmission dynamics in island net-works. In this study, we characterised the distribution and activity patterns of BN and LN, and assessed species-related and inter-annual variation in AIV shedding and seroprevalence. Our findings demonstrate the very diverse dispersive behaviours of BN and LN, and the high connectivity between islands generated by post-breeding migrations. Indeed, although based on a relatively limited number of tracked birds and a single breeding site, we detected a potential connectivity with ten different roosting islands. Epidemiological data further support that these species are major hosts for AIV on tropical islands. At the scale of the Western Indian Ocean, BN and LN migratory behaviours could therefore have major implications for AIV transmission between islands.

Specificity in the ecology of terns might account for the variation in virus infection and transmission between closely related species. For instance, species-related differences in life history such as colonial nesting, social behaviour, migration and foraging characteristics could affect opportunities for virus transmission. The extensive post-breeding migration of ST (average travelled distance > 50 000 km), together with their infrequent contact with seawater and time spent on the ground [35], may for instance limit infection opportunities. This species also never roost on islands during the post-breeding migration and may therefore have a more limited role than BN and LN in AIV interisland transmission. For both BN and LN, our data showed that tracked birds either stayed in the vicinity of their breeding ground, or moved to and roosted on another island, without major differences in the proportion of resident *vs* migrant birds between species. However, given the significant difference of AIV seroprevalence between BN and LN, other factors than migratory and activity patterns are likely to be involved in virus transmission.

Differences in the diversity and location of roosting sites selected by BN and LN may also account for the variation of AIV exposure. Indeed, one may hypothesize that virus maintenance and transmission could be heterogeneous at the scale of the Western Indian ocean (i.e. between islands), therefore generating a higher proportion of seropositive birds for species migrating to AIV circulation areas. Our tracking data did not reveal species-related variation in bird migration but rather individual behaviours in the selection of roosting islands. Factors involved in the selection of roosting sites include nearby predictable availability of prey, which are often associated with the proximity of larger sub-surface predators [36] and oceanographic features including upwellings and countercurrents [37]. Conversely, seabirds might avoid regions prone to adverse weather, especially storms [38]. In order to identify the links between tern migratory patterns and AIV transmission, future studies will have to focus on the drivers and repeatability of migratory decision making at the individual level, as well as inter-annual variation in the selection of roosting sites and associated infectious and immune status of migrant birds.

Beyond knowledge on tropical tern migrations, current understanding of AIV transmission dynamics in tropical seabirds remains limited. Although the seasonal increase of the number of fledglings at the end of the breeding season could drive virus transmission in seabird colonies [39], mechanisms involved in the inter-annual maintenance of these viruses remain to be identified. In this study, none of the samples tested positive for AIV RNA. This may be because sampling was conducted mostly on adult birds that already had mounted AIV-specific immunity, but also because sampling occurred mostly before hatching and the incoming of immunologically naïve chicks into the population. Seroconversion was detected in recaptured adult BN, suggesting that infection occurred during the course of the study. Differences in seroconversion patterns could be associated with AIV subtype-specific variation in the long-term immune response and protection, as demonstrated for H13 and H16 [40]. Longitudinal studies are needed to precisely assess the temporal variation and drivers of AIV transmission on tropical islands. Because most seabird species leave their breeding colonies after chick fledging, virus detection after the breeding season is rarely feasible. Further epidemiological studies on resident BN and LN could, however, provide information on the ecological drivers of viral infection but also on the timing of virus introduction and transmission dynamics in highly isolated seabird populations.

Seabirds are highly sensitive to environmental changes and significant modifications of their biology in response to climatic [40] and anthropogenic changes, such as habitat modifications and alien predator introductions [41], have been described. Seabird numbers have decline by 70% over the past 60 years, with the highest decreased reported for terns (86%; [42]). Terns are impacted by environmental changes in several distinct ways, affecting both marine and coastal habitats. Future studies focusing on AIV epidemiology on tropical oceanic islands will also need to consider the potential cascade effects of environmental changes, population decline and individual stress on virus transmission dynamics.

## Acknowledgments

We are very grateful to the Savy family for their warm hospitality and support in the field work on Bird Island. We also thank Brigitte Pool, Graham Govinden, and Léon Biscornet, for arranging sample conservation at the Victoria Hospital.

## Author contributions

Study design: CLe, MLC, AJ. Funding acquisition: CLe, MLC. Field work: CLe, MD, CF, CLa, AJ. Serology and molecular analyses: CLe. Molecular sexing: SB, LH. Data analyses: CLe, SB, AJ. Manuscript preparation: CLe, AJ. All authors edited, read, and approved the final manuscript.

---

**Supplementary Table 1.** Origin of serum samples.

| Year | Species | Number of positive sera | Number of tested sera | Reference |
| --- | --- | --- | --- | --- |
| 2012 | *Anous stolidus* | 11 | 31 | [2] |
|  | *Anous tenuirostris* | 23 | 50 |  |
|  | *Onychoprion fuscatus* | 15 | 207 |  |
| 2013 | *Anous stolidus* | 28 | 90 | [2] |
|  | *Anous tenuirostris* | 51 | 90 |  |
|  | *Onychoprion fuscatus* | 10 | 100 |  |
| 2014 | *Anous stolidus* | 41 | 151<sup>1</sup> | This study |
|  | *Anous tenuirostris* | 97<sup>3</sup> | 171<sup>2</sup> |  |
|  | *Onychoprion fuscatus* | 9 | 107 |  |
| 2015 | *Anous stolidus* | 27 | 133 | This study |
|  | *Anous tenuirostris* | 58 | 101 |  |
|  | *Onychoprion fuscatus* | 3 | 53 |  |

<sup>1</sup> Samples collected from 144 adults, four juveniles and three chicks.
<sup>2</sup> Samples collected from 141 adults and 30 chicks.
<sup>3</sup> Seven of the 97 positive samples were collected in chicks.

**Supplementary Table 2.**
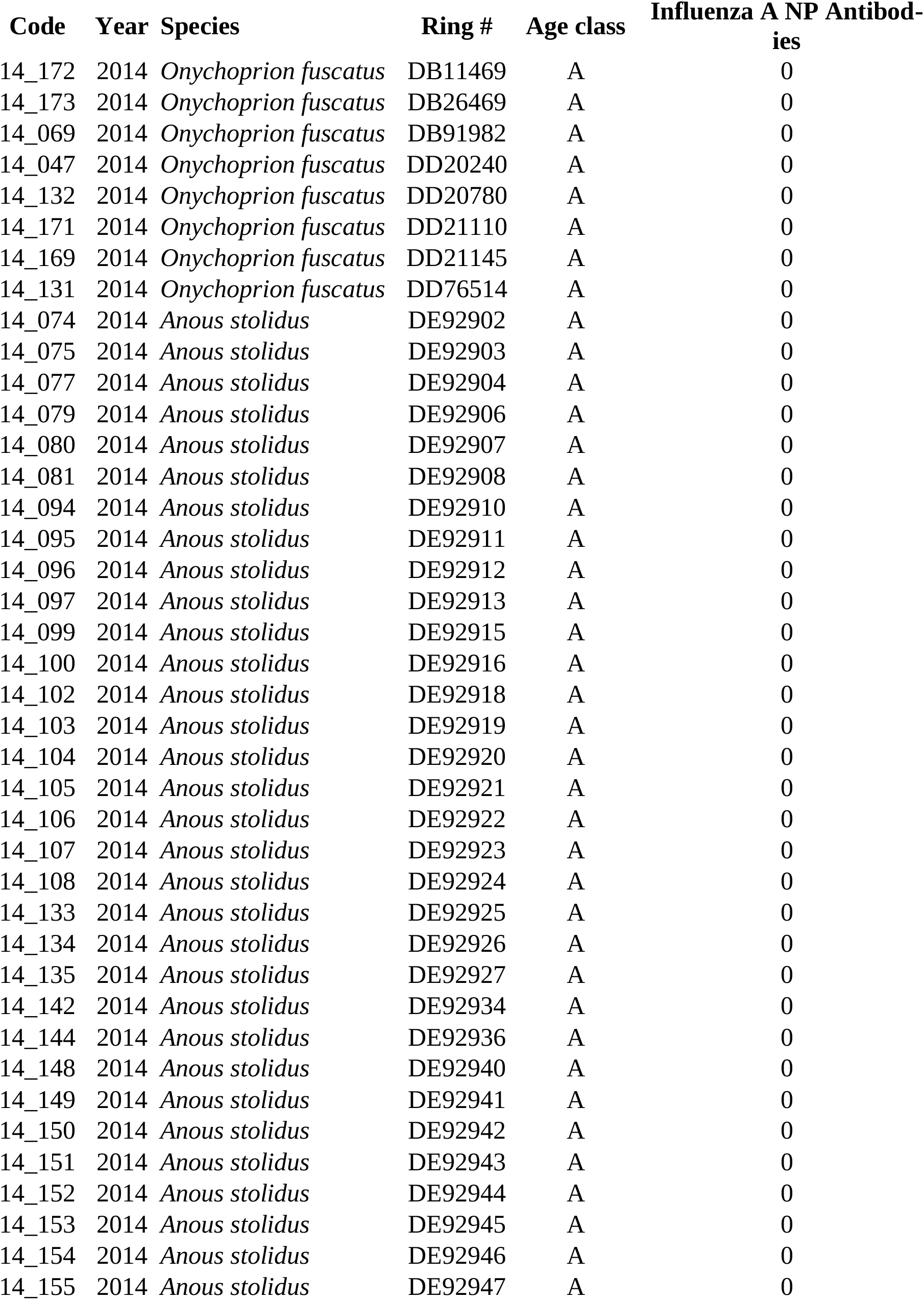
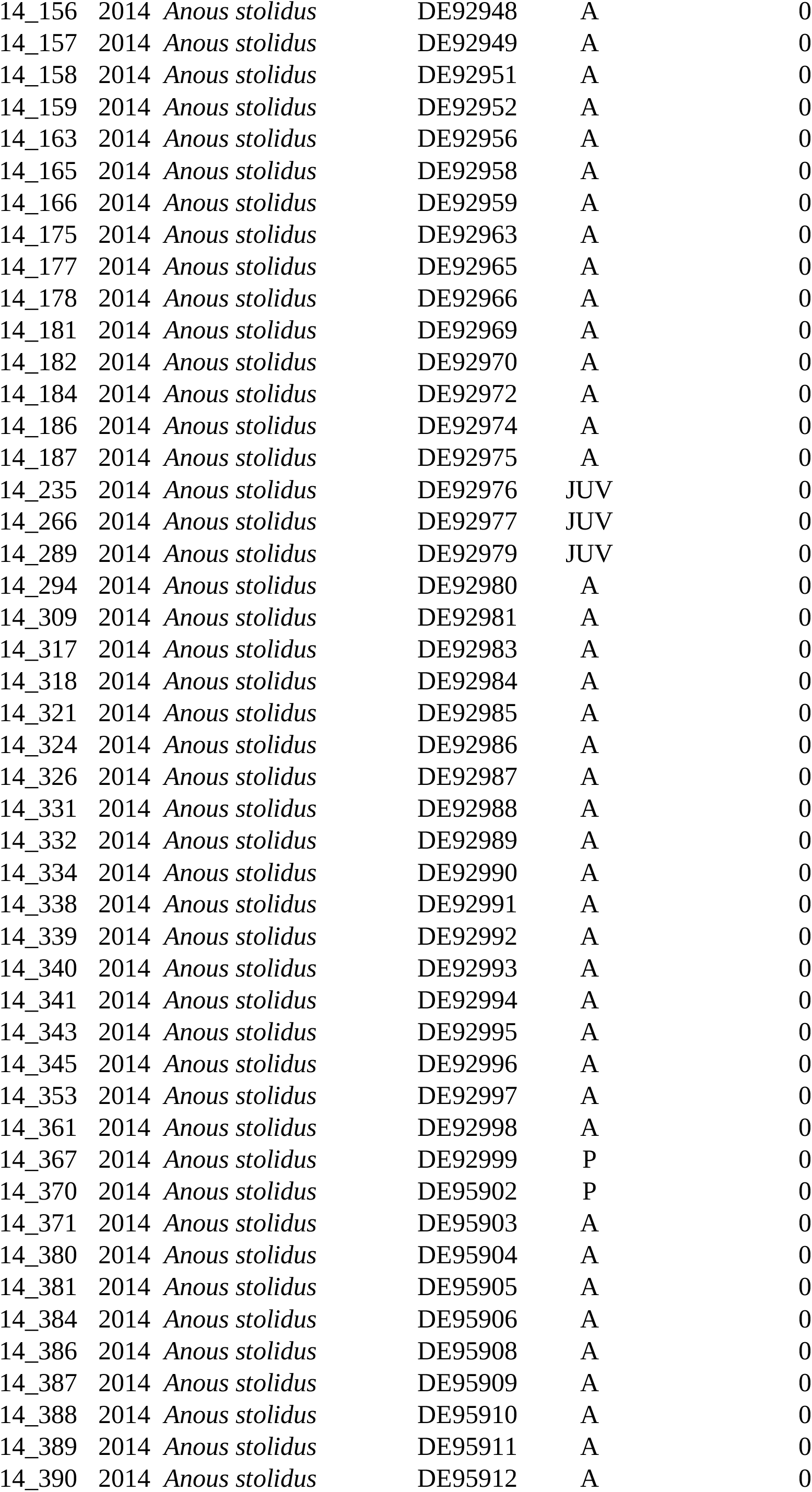
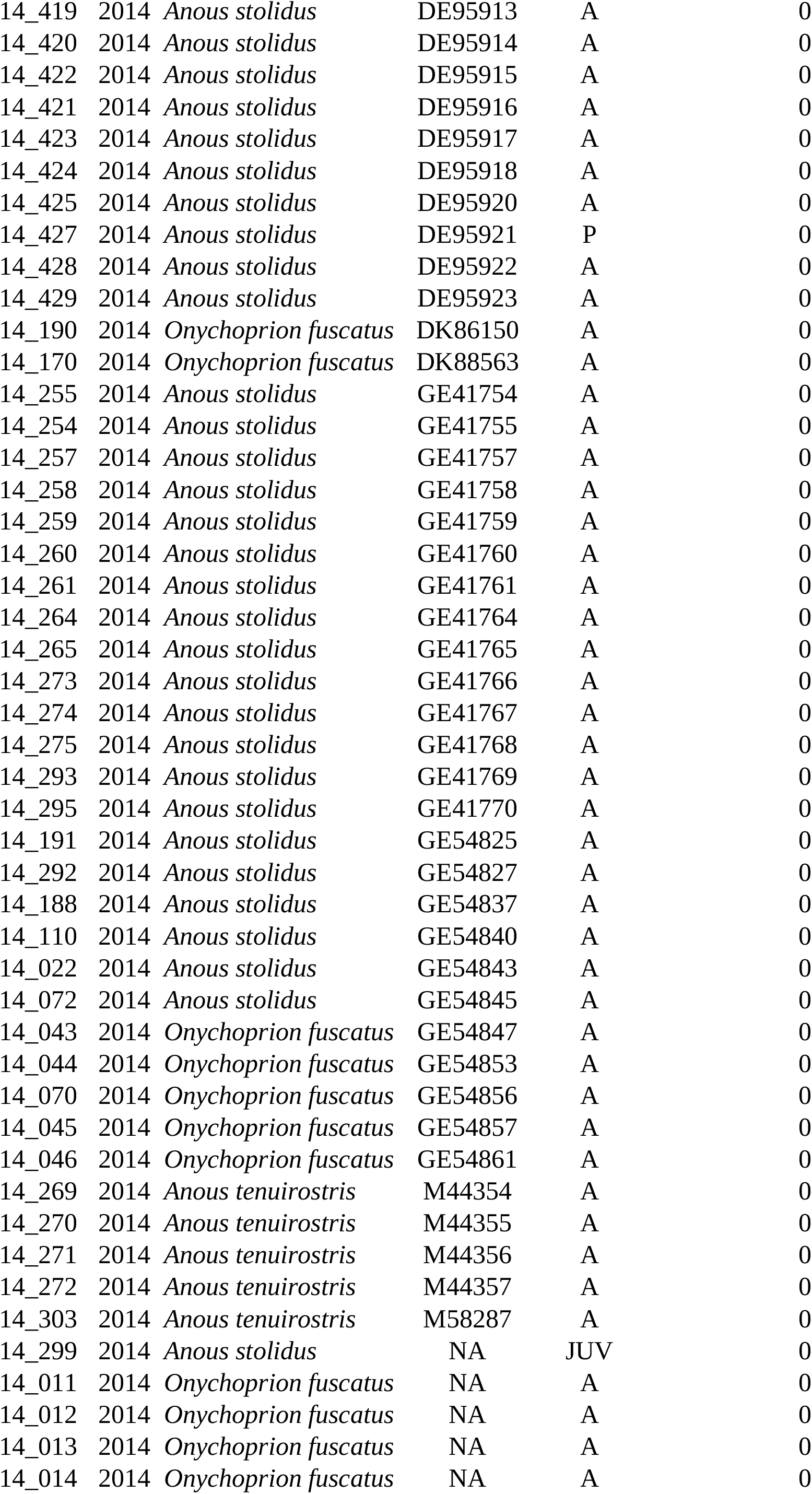
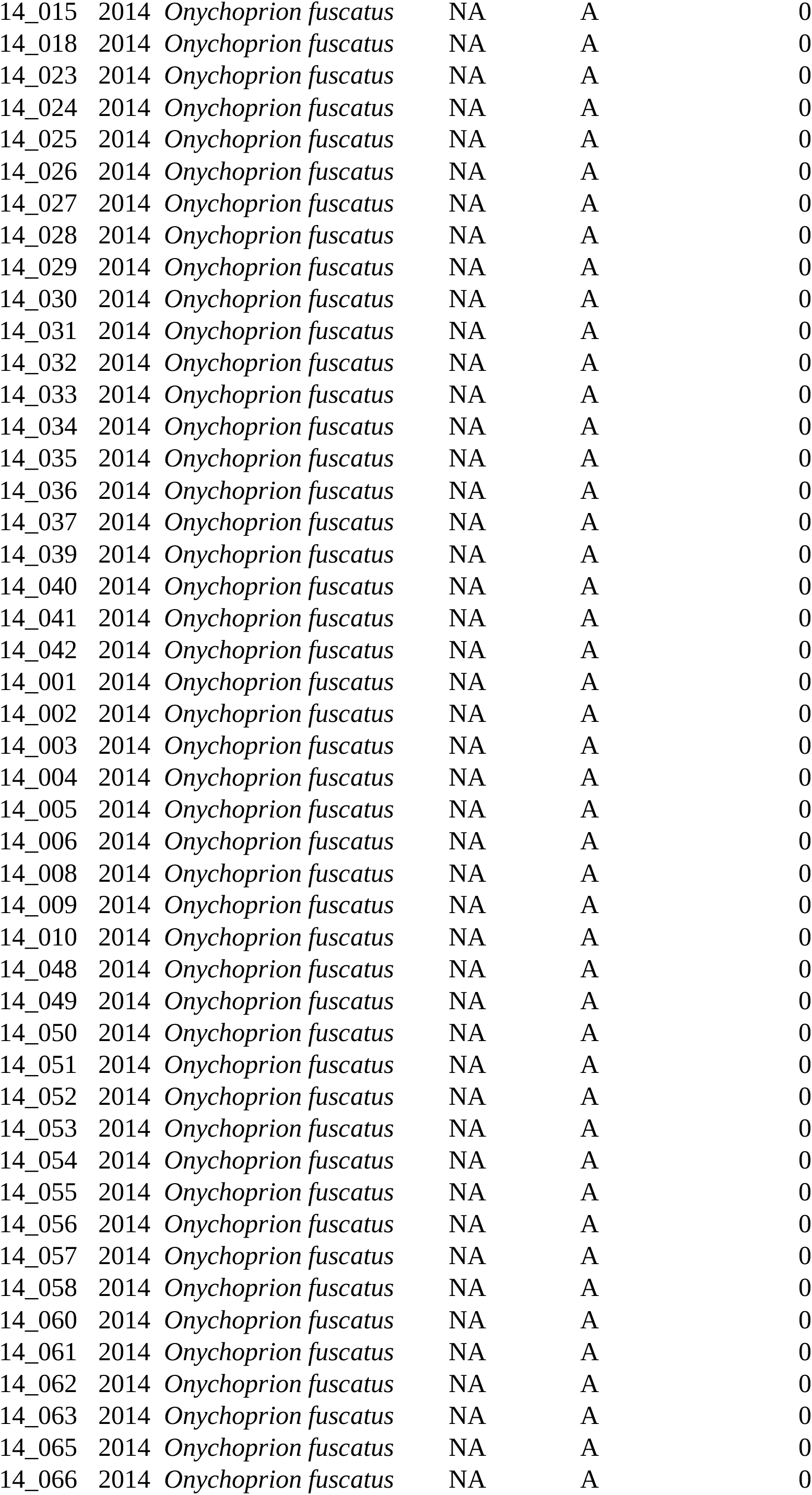
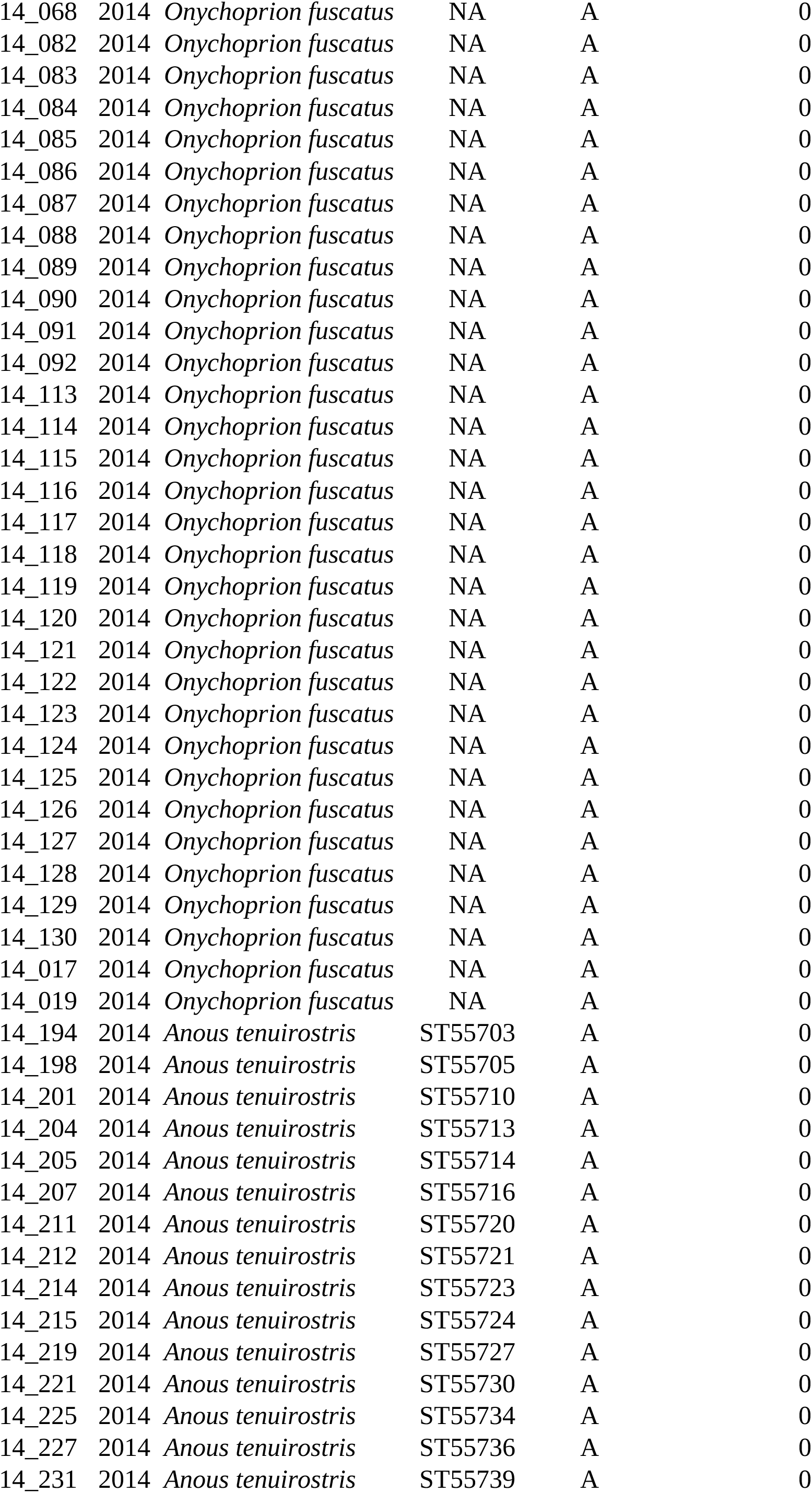
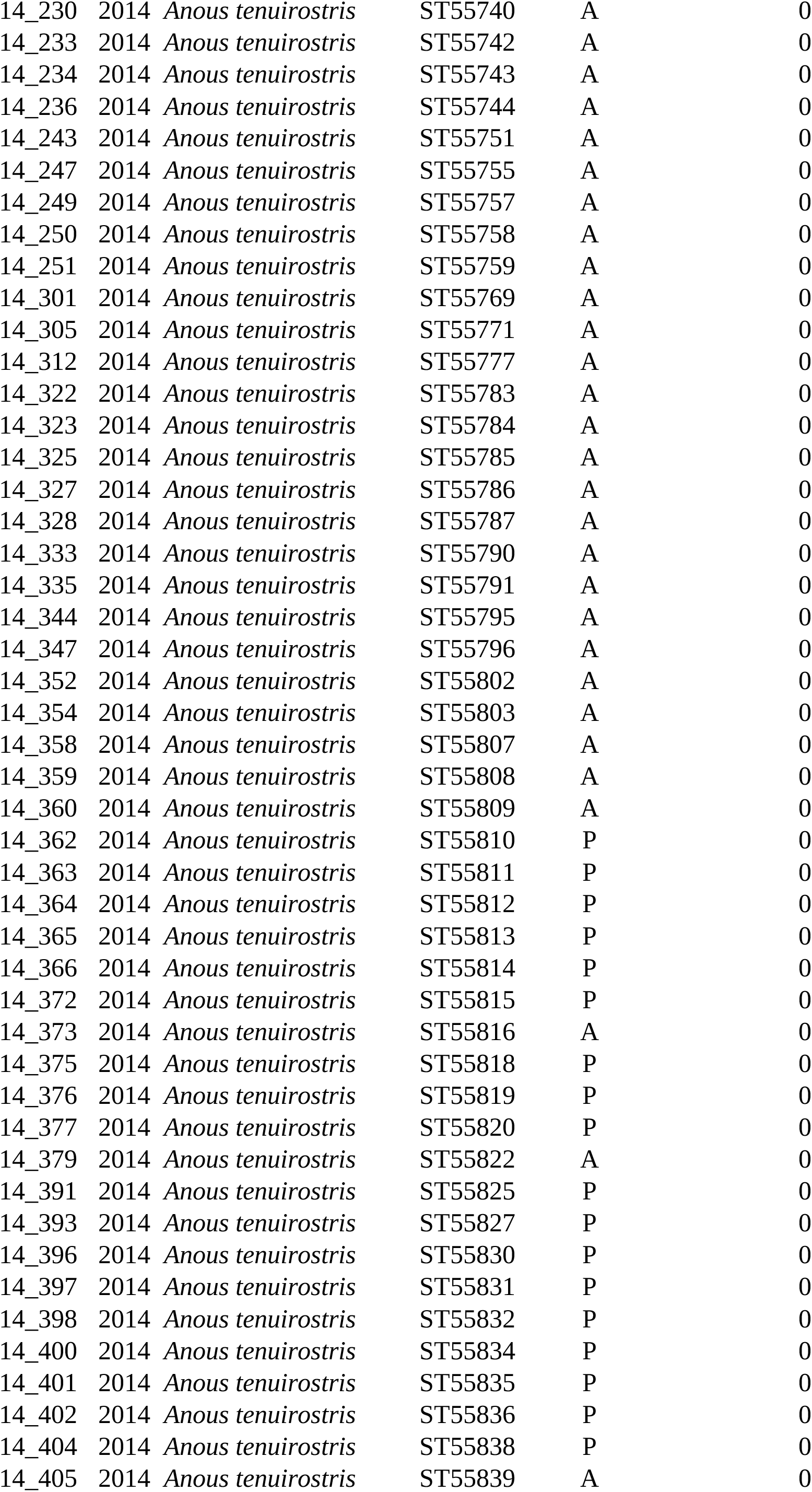
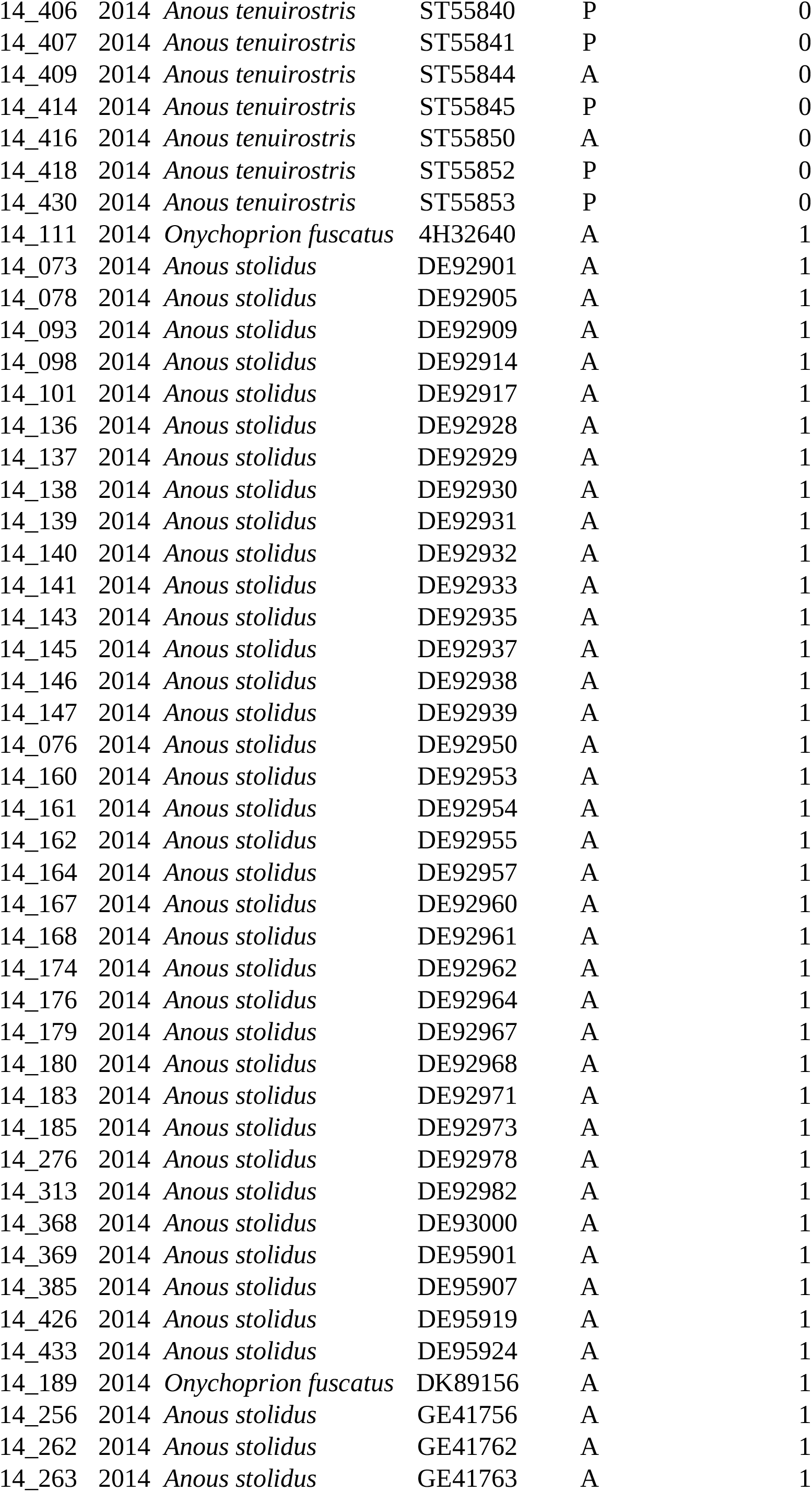
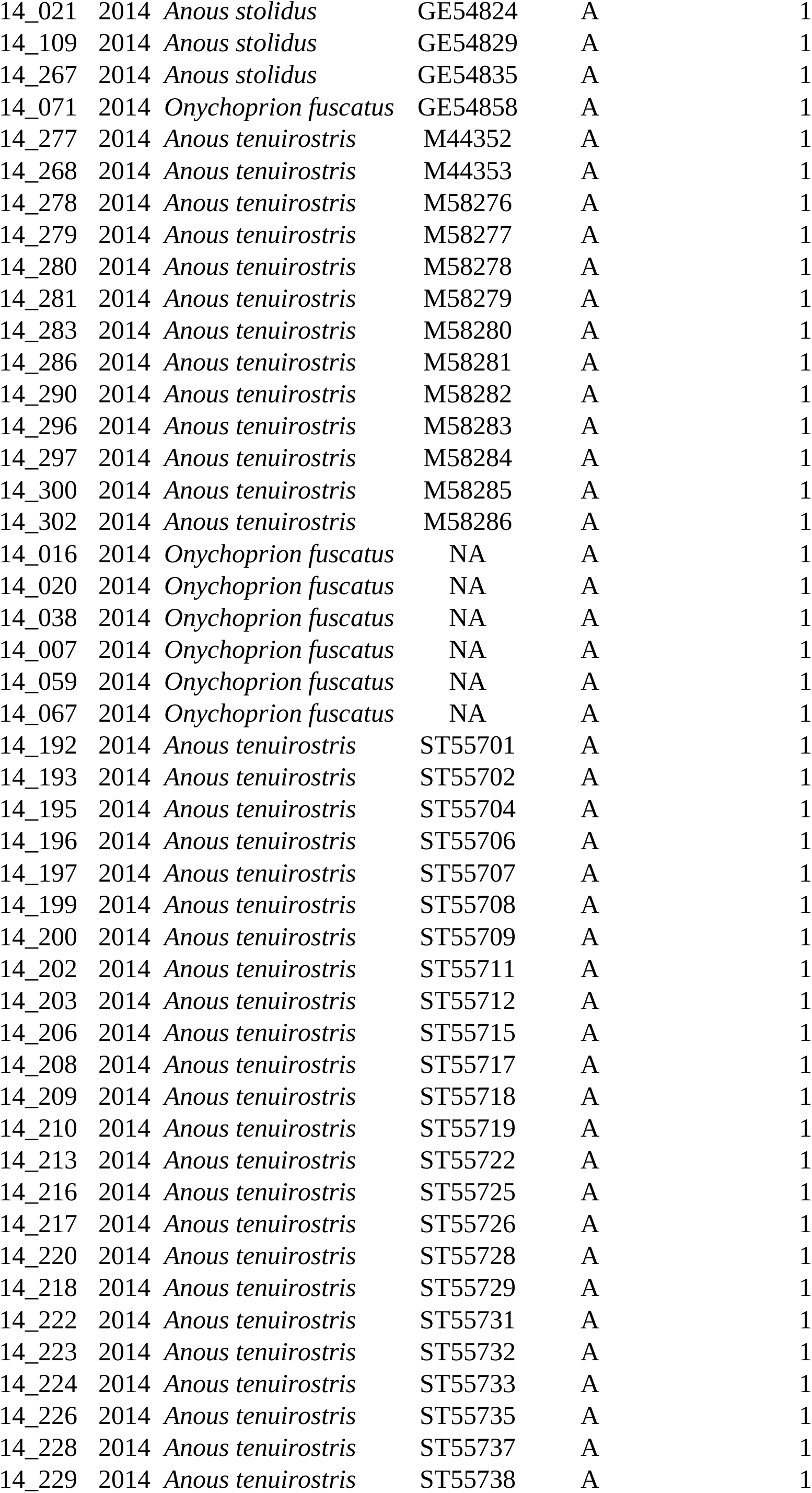
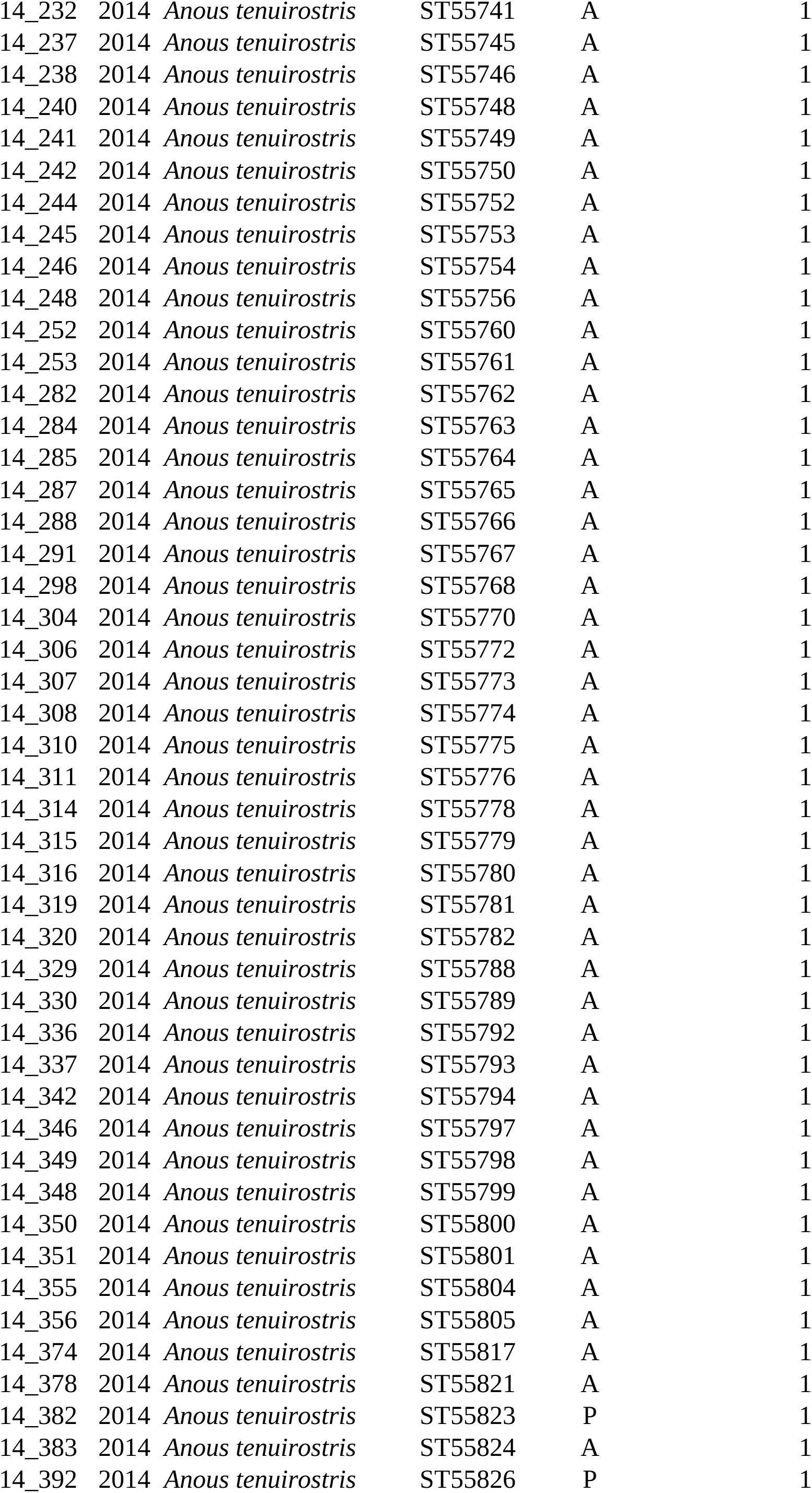
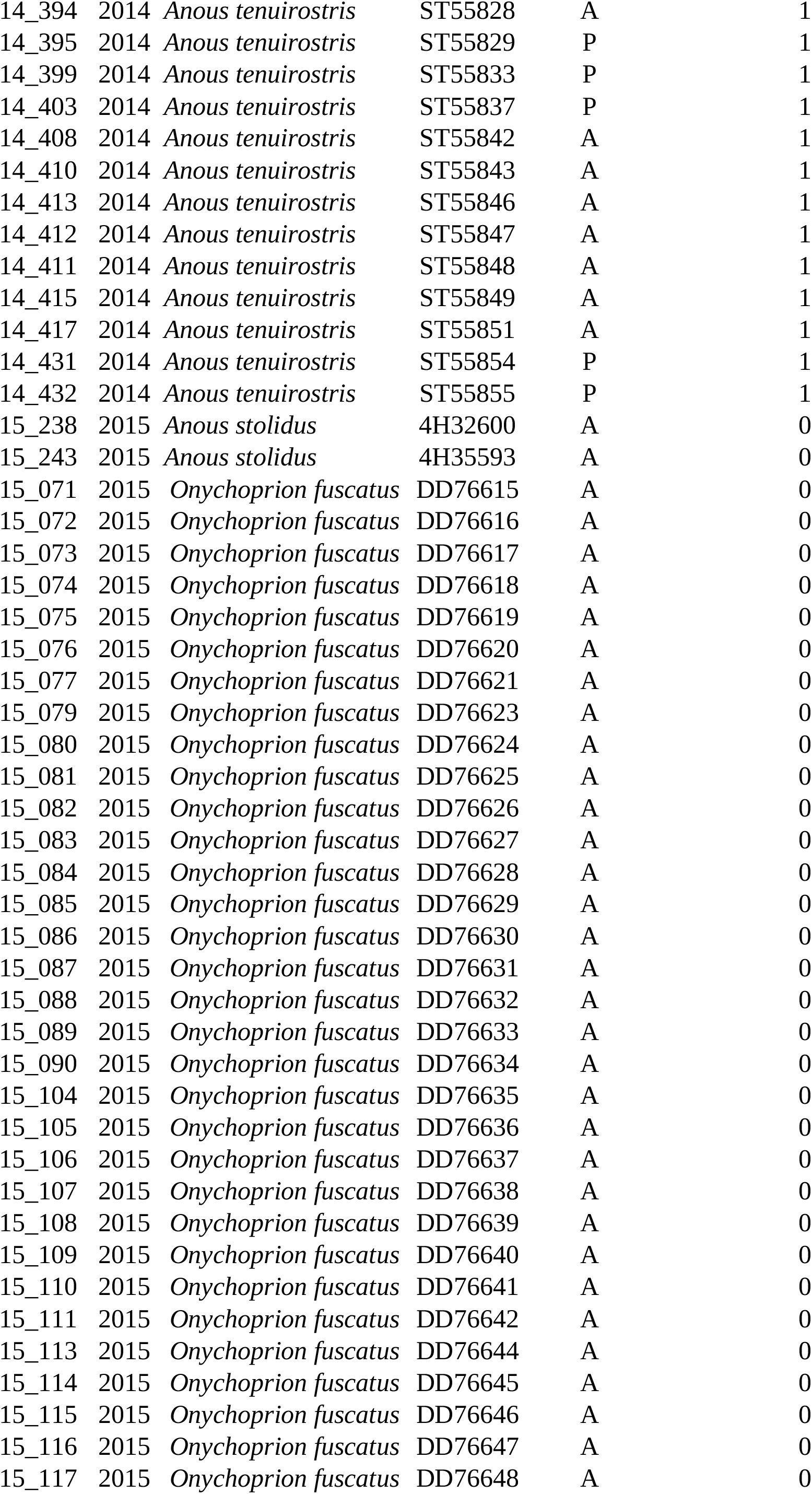
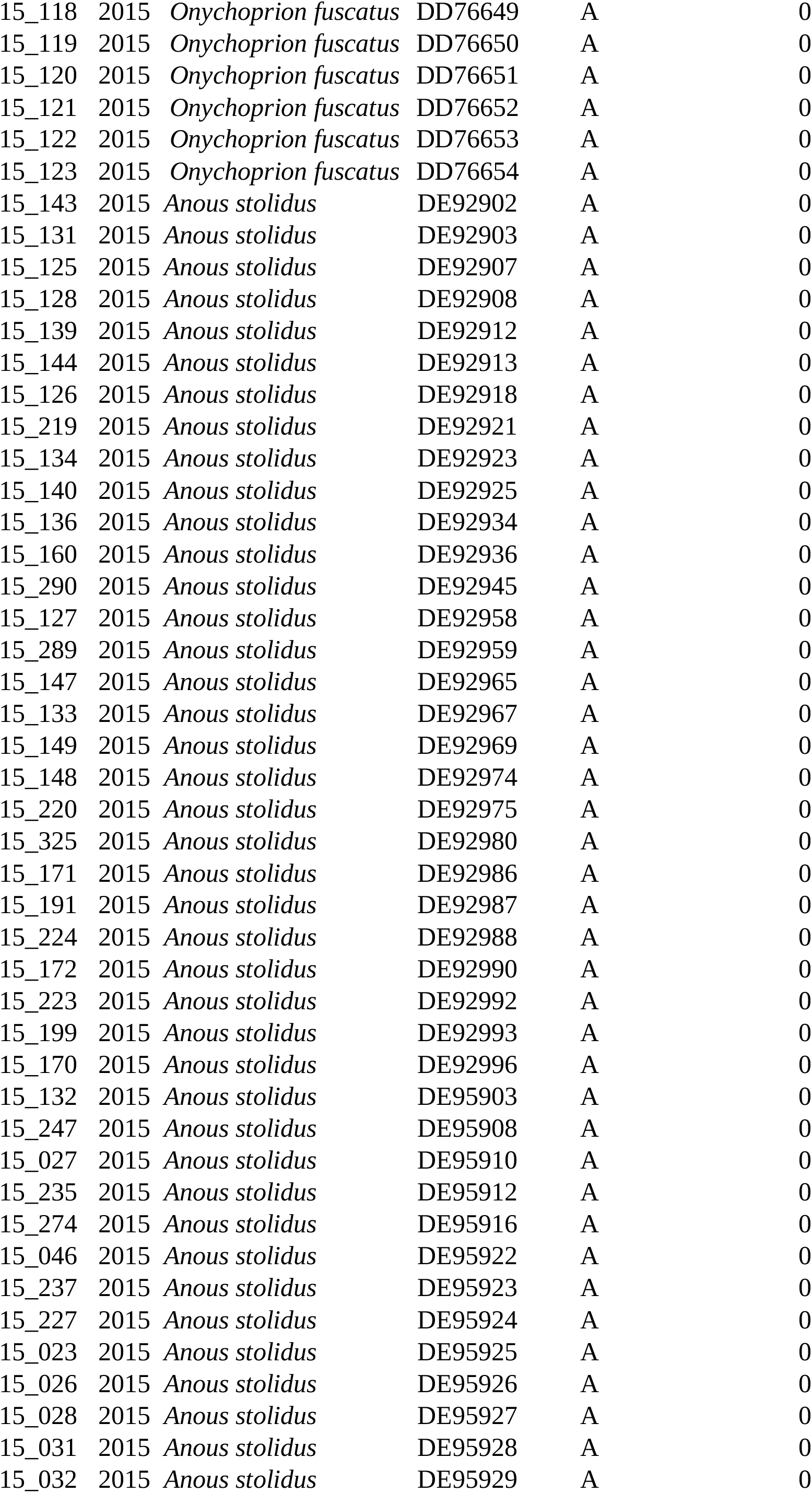
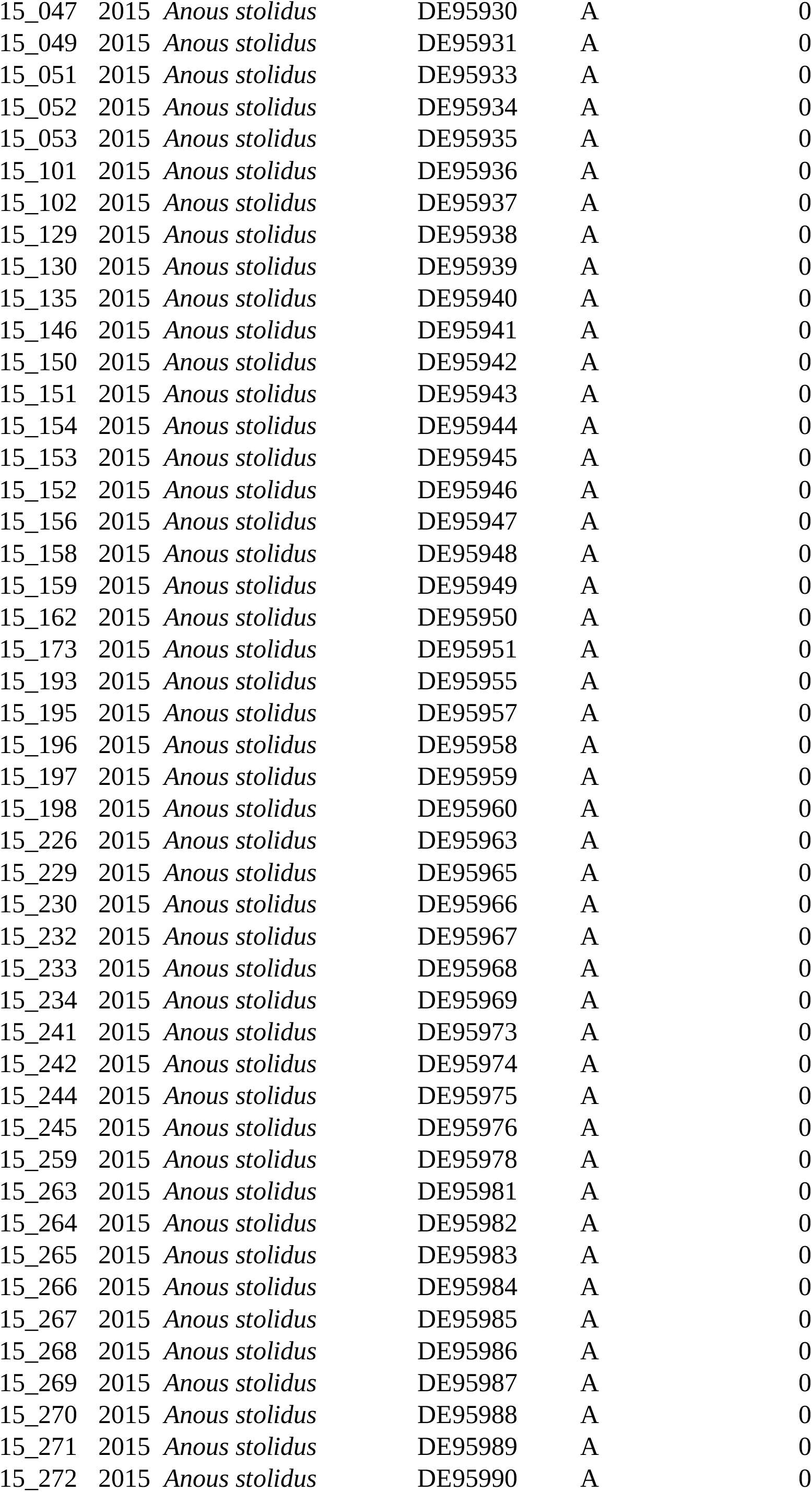
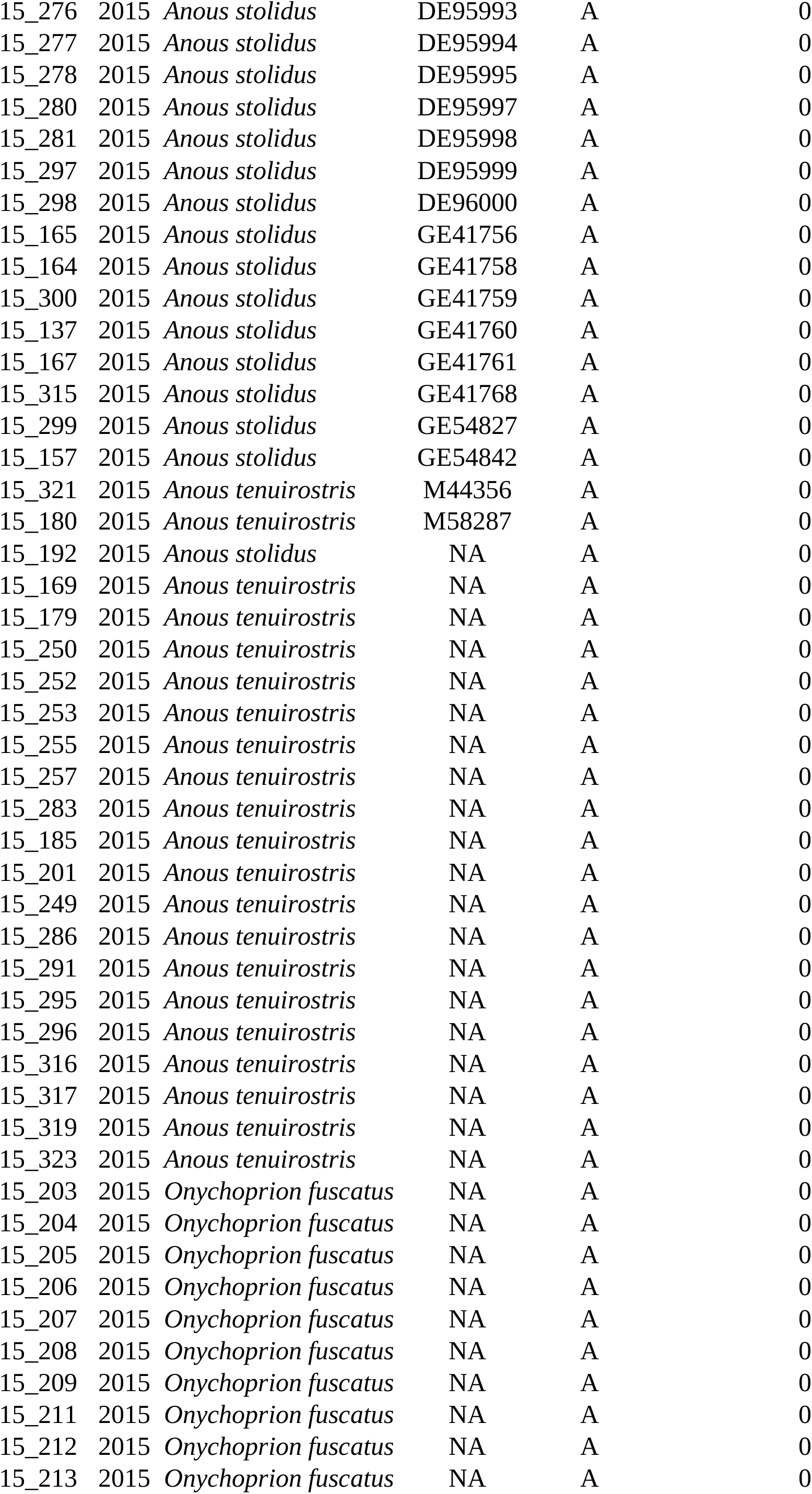
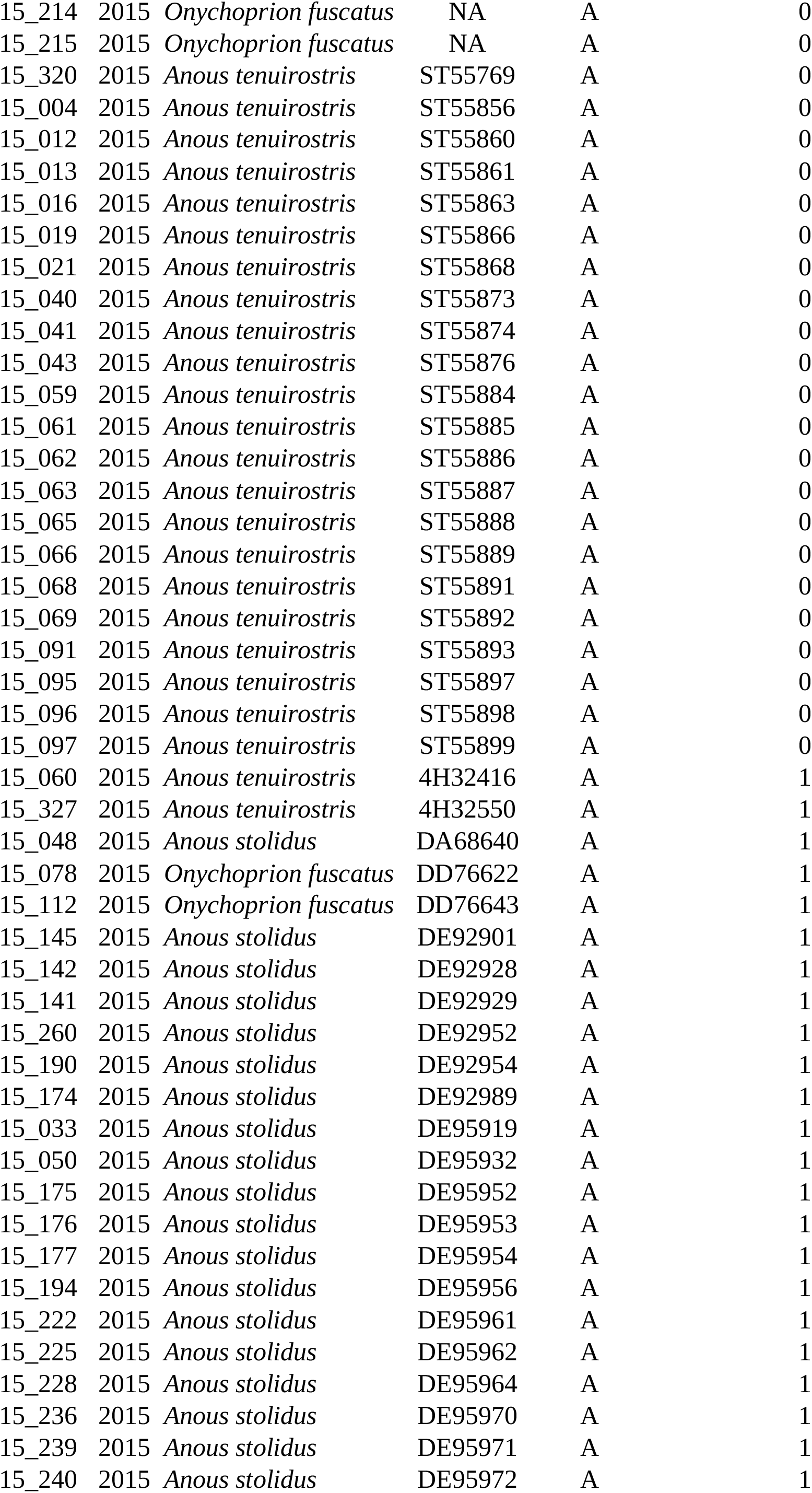
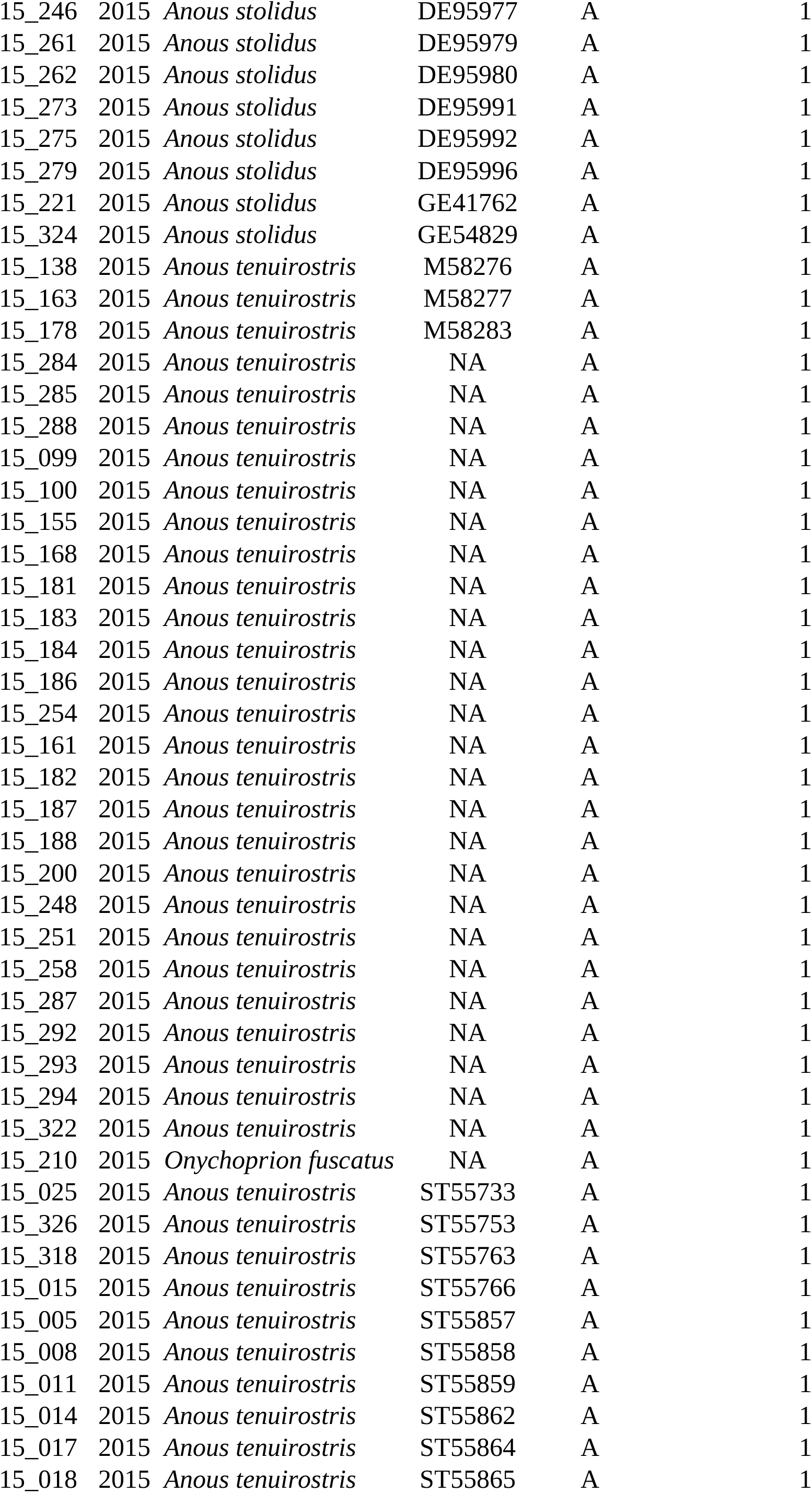
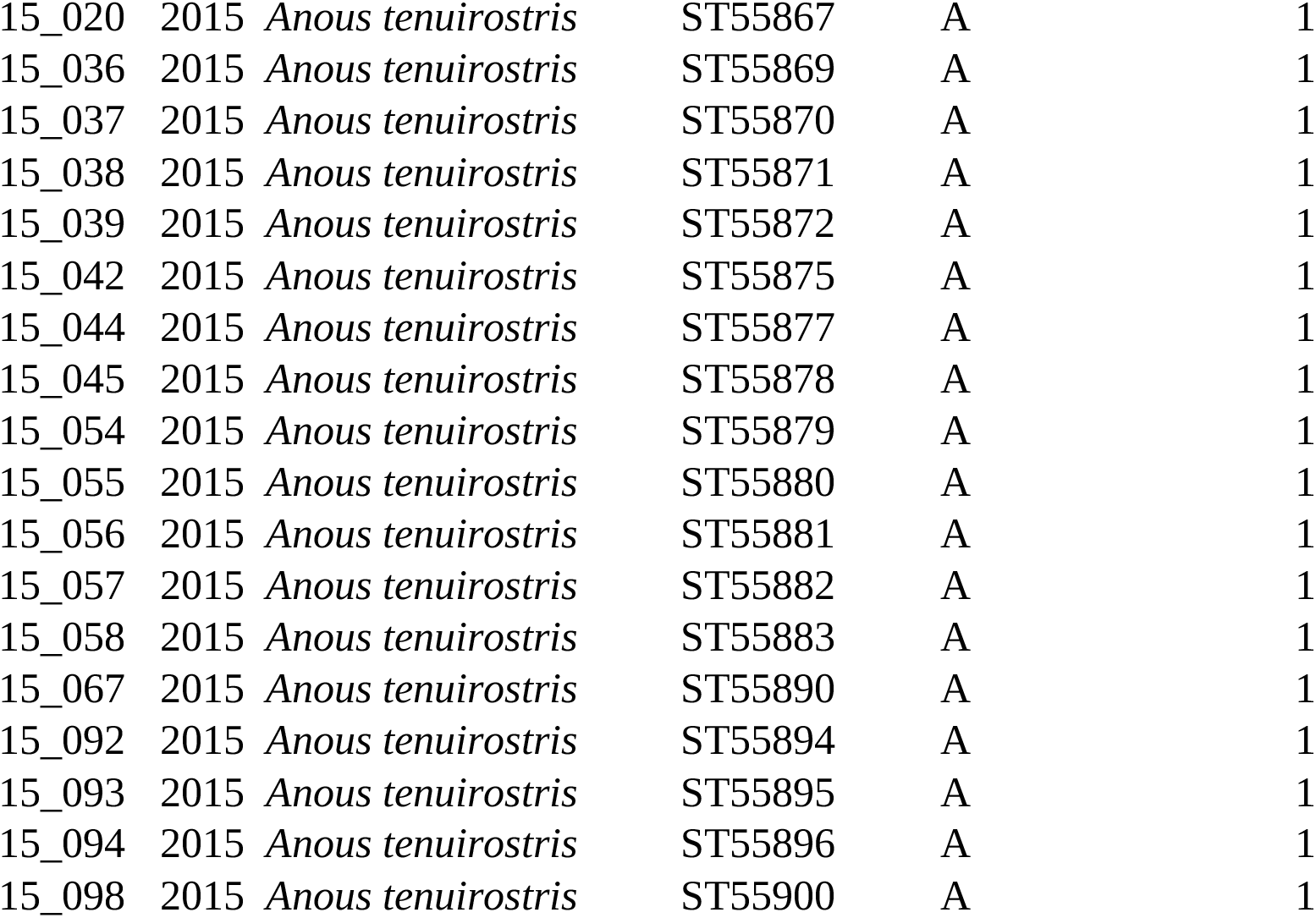
Detection of avian influenza virus nucleoprotein (NP) specific antibodies in 2014 and 2015.

**Supplementary Table 3.**
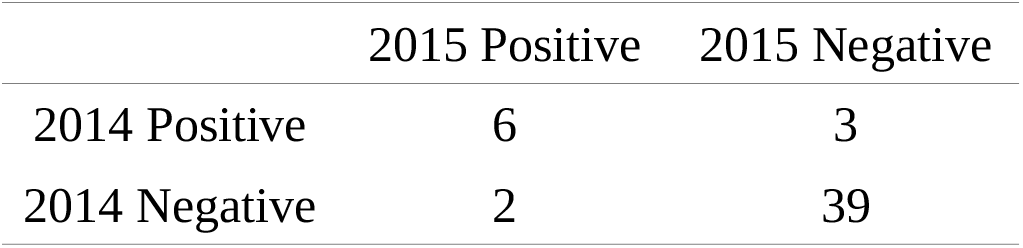
Detection of avian influenza virus nucleoprotein specific antibodies in Brown noddies (*Anous stolidus*), in 2014 ringed birds recaptured in 2015.

|  | 2015 Positive | 2015 Negative |
| --- | --- | --- |
| 2014 Positive | 6 | 3 |
| 2014 Negative | 2 | 39 |

